# Stimulation-Evoked Effective Connectivity (SEEC): An in-vivo approach for defining mesoscale corticocortical connectivity

**DOI:** 10.1101/2022.03.03.482925

**Authors:** David T. Bundy, Scott Barbay, Heather M. Hudson, Shawn B. Frost, Randolph J. Nudo, David J. Guggenmos

## Abstract

**Background:** Cortical electrical stimulation has been a versatile technique for examining the structure and function of cortical regions as well as for implementing novel therapies. While electrical stimulation has been used to examine the local spread of neural activity, it may also enable longitudinal examination of mesoscale interregional connectivity. Recent studies have used focal intracortical microstimulation (ICMS) with optical imaging to show cross-region spread of neural activity, but techniques are limited to utilizing hemodynamic responses within anesthetized preparations.

**Objective:** Here, we sought to use ICMS in conjunction with recordings of multi-unit action potentials to assess the mesoscale effective connectivity within sensorimotor cortex.

**Methods:** Neural recordings were made from multielectrode arrays placed into sensory, motor, and premotor regions during surgical experiments in three squirrel monkeys. During each recording, single-pulse ICMS was repeatably delivered to a single region. Mesoscale effective connectivity was calculated from ICMS-evoked changes in multi-unit firing.

**Results:** Multi-unit action potentials were able to be detected on the order of 1 ms after each ICMS pulse. Across sensorimotor regions, short-latency (< 2.5 ms) ICMS-evoked neural activity strongly correlated with known anatomic connections. Additionally, ICMS-evoked responses remained stable across the experimental period, despite small changes in electrode locations and anesthetic state.

**Conclusions:** These results show that monitoring ICMS-evoked neural activity, in a technique we refer to as Stimulation-Evoked Effective Connectivity (SEEC), is a viable way to longitudinally assess effective connectivity enabling studies comparing the time course of connectivity changes with the time course of changes in behavioral function.

**Highlights:**

- Short-latency neural responses to ICMS were evaluated in multiple cortical regions.
- Neural responses strongly correlated with known anatomical connections.
- Stimulation-evoked neural responses were maintained across repeated tests.
- ICMS-evoked activity can show longitudinal changes in effective connectivity.

## Introduction

Electrical stimulation of the brain is a well-established clinical and scientific tool. Clinically, stimulation can be used to modify the excitability of neurons (1, 2), map functional brain regions (3), and evoke artificial sensory percepts (4, 5). In the laboratory setting, short-duration microampere-current pulses can be delivered through small microelectrodes implanted into the cortex using a technique known as intracortical microstimulation (ICMS). ICMS modulates neural activity within a focal region of the brain and has helped to elucidate the structure and function of multiple brain regions. ICMS in awake behaving animals can modulate behavior, which gives insight into the functional roles of brain regions (6). ICMS has also been used to examine plasticity in motor output maps following behavioral training and post-stroke recovery (7-11). While there are numerous examples of the effects of ICMS on sensory perception, muscle activation and neuroplasticity mechanisms, and the mode of somatic activation continues to be explored (12), few studies have investigated the transsynaptic effects of ICMS, especially in remote cortical regions.

The output properties of ICMS are dependent on both the direct and indirect effects on neurons and neural circuits. Stimulation directly induces voltage-gated ion channels in neurons to open, with axons activated at lower currents than cell bodies (12-15). However, the specific neural components activated depend on the excitability of the neural component, distance from the electrode, stimulus amplitude, and stimulus duration (14, 15). Even at low amplitudes, stimulation can evoke transsynaptic activity (16), which underlies the potential to use stimulation to examine corticocortical connectivity.

Corticocortical connectivity analyses typically rely on examining the pattern of fiber degeneration following a lesion (17) or on examining the transport of injected anatomical tracers (18-21). These methods have revealed extensive interareal connectivity, but they are limited to injecting tracers into a few regions at a single time point. Because changes in connectivity are associated with neural injuries and disorders, this cross-sectional approach is a significant limitation in interpreting the role of structural reorganization in functional recovery. An alternative to these anatomical approaches is to measure evoked neural activity following ICMS. Recently, this method has been used with intrinsic optical imaging and functional magnetic resonance imaging (fMRI) to demonstrate intra- and interregional connectivity (22-26). However, because these modalities measure the hemodynamic response, it is difficult to separate synaptic activity from neural firing in order to isolate direct anatomical projections as is possible with short latency responses (26).

We propose a novel method to examine mesoscale corticocortical connectivity using direct recordings of multi-unit neural activity to measure ICMS-evoked neural activity. When combined with an algorithm to remove stimulus artifact, we were able to examine short-latency neural responses to electrical stimuli on the order of 1 ms after a stimulus pulse. Importantly, this allows us to identify direct connections by differentiating short-latency action potentials from either polysynaptic action potentials or sub-threshold synaptic potentials.

## Methods

### Experimental Design

We performed intraoperative recordings in 3 adult (12 – 17 years old) male squirrel monkeys (*Saimiri sciureus*), a New World monkey with a relatively lissencephalic cortex, allowing visualization of sensorimotor regions on the surface of the brain. All procedures were approved by the Institutional Animal Care and Use Committee at the University of Kansas Medical Center in compliance with *The Guide for the Care and Use of Laboratory Animals* (27). At the conclusion of these experiments, as part of the same surgery, monkeys were utilized for a secondary terminal study (28).

### Motor Mapping

Surgical procedures have been described previously (10). Following a craniectomy, motor and premotor cortices were mapped using ICMS under ketamine anesthesia. A photograph of the exposed cortex was taken to enable co-registration of ICMS-evoked responses with cortical landmarks. For initial neurophysiological identification of motor areas, a microelectrode was made from a glass micropipette tapered to a fine tip, filled with 3.5 M NaCl, and inserted into cortex to a depth of 1750 μm (Layer 5). Pseudo-biphasic stimulus pulses were delivered as described previously (cathode leading, 200 μs/phase, 13 pulse train at 350 Hz, repeated once per second) (10). Stimulation current was increased slowly until a visible movement was elicited on at least 50% of stimuli or a maximum current of 30 μA was reached with no response observed. Electrode penetrations were made at a spatial resolution of ∼500 μm with finer resolution used as needed to delineate the borders between regions. The primary motor cortex (M1-DFL), ventral premotor cortex (PMv-DFL), and dorsal premotor cortex (PMd-DFL) distal forelimb regions were identified by defining boundaries between sites where stimulation evoked digit or wrist movements and sites with more proximal elbow, shoulder, trunk or face movements (7, 9-11). In two monkeys, the face region (PM-face) rostral to M1-DFL was also identified.

### Somatosensory Mapping

To determine boundaries between somatosensory regions (areas 3a, 3b, 1, and 2/5) a 16-contact linear multi-electrode array (Neuronexus, Ann Arbor, MI) was inserted into cortex to record multi-unit activity within a cortical column. Signals were filtered, amplified, and played on a speaker to monitor modulation in multi-unit firing. Cutaneous receptive fields were defined using Semmes-Weinstein monofilaments, and deep receptive fields were determined using high-threshold manual stimulation and joint manipulation. Area 3a was characterized by multi-digit cutaneous receptive fields, and areas 3b and 1 were characterized by small cutaneous receptive fields that mirrored each other with distal to proximal hand sites arranged in rostral-to-caudal (area 3b) and caudal-to-rostral (area 1) orientations (29). Caudal to area 1, sites had multi-digit receptive fields sensitive to proprioceptive stimuli. As the presence of area 2 in squirrel monkeys is uncertain (30, 31), sites caudal to area 1 were classified as area 2/5.

### Neural Recordings

Neural activity was recorded from the identified regions during single-pulse ICMS. All recordings and stimulation were made with an Intan RHS Stim/Record system (Intan Technologies, Los Angeles, CA). Three 32-channel microelectrode arrays (4 shanks x 8 sites per shank, 703 μm^2^ site area, 100 μm site spacing, 400 μm shank spacing) were used. One array was placed in S1 (targeting areas 3a, 3b, 1, or 2/5), a second was placed in M1-DFL, and a final array was placed in premotor cortex (targeting PMv-DFL, PMd-DFL, or PM-face). Microelectrode arrays were inserted to depths of 1750 μm (M1 and PM) and 1500 μm (S1). In monkey 1 two arrays had four sites activated to reduce electrode impedance, allowing for stimulation and recording. In monkeys 2 and 3, all three arrays were activated for stimulation and recording. After placing the arrays, recordings were made with 1,000 ICMS pulses delivered through one of the activated microelectrode sites in one array while neural activity was recorded at 30 kHz. Each ICMS stimulus consisted of a single, cathodal-leading biphasic pulse (200 μs/phase, 50 μA); ICMS stimuli were repeated at the rate of 5 Hz, allowing ICMS-evoked neural activity to be recorded for up to 200 ms post-stimulus. The site with the lowest impedance on each array was chosen for stimulation. After recordings were made stimulating through each array, all three arrays were removed from the cortex, either the sensory or premotor array was moved to a new location in a pseudo-random order, and the arrays were reinserted. This was repeated until each combination of sensory and premotor regions was tested. Except for PM-face, each region with an activated array was stimulated.

### Neural Data Processing

ICMS-evoked neural activity was analyzed offline. Following the stimulus pulse (200 μs/phase), the post-stimulus electrical artifact consisted of a brief period of amplifier saturation with a slow fall-off lasting over 10 ms (Fig 1A). For each channel, this period of saturation was determined quantitatively to set the absolute blanking period from 0.5 ms before the stimulus pulse to 0.25 ms after amplifier saturation ended. Data within this blanking period were set to 0. The remaining portion of the stimulation artifact was removed using a sliding polynomial fit (32). An 8^th^ order polynomial was fit to the signal following each stimulus pulse using a 6 ms sliding window. The fit data was then subtracted from the recorded data allowing action potentials to be detected ∼1 ms after each stimulus pulse. Periods with artifact were excluded using a nonlinear energy operator and data was high-pass filtered with a single-pole IIR high-pass filter with a cutoff frequency of 500 Hz. Finally, multi-unit action potentials were detected using a threshold detector set at -3.5x the RMS voltage for each channel (Fig. 1B).

**Figure 1:**
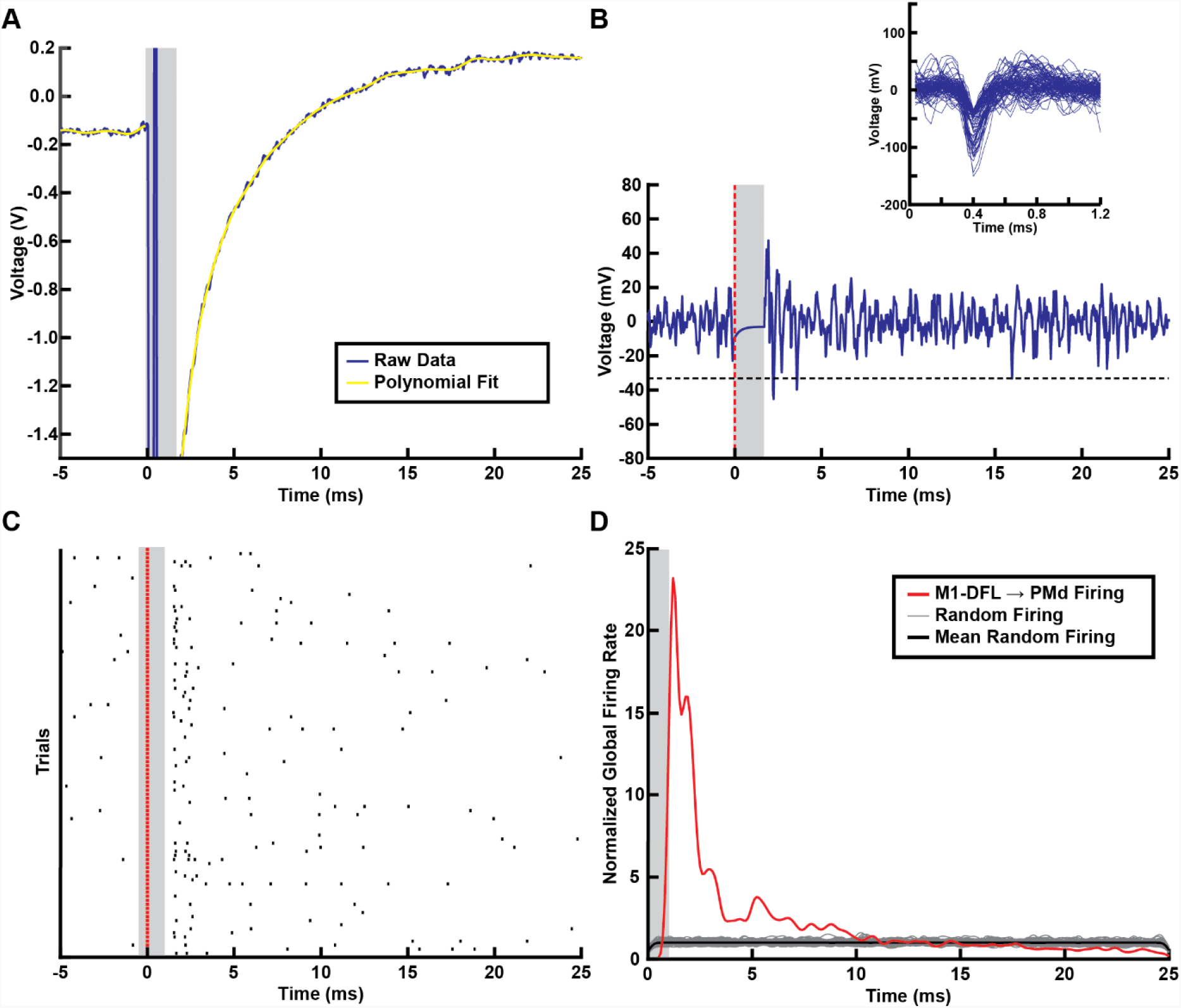
Stimulus artifact correction. **A**. Single-pulse ICMS stimulation produced an initial large amplitude artifact that with a slow return to baseline. The unusable period of the data (∼1 ms post-stimulus) in which the stimulation was ongoing, or the amplifiers were saturated was automatically detected and set to zero. A sliding polynomial fit was used to detect the remaining stimulus artifact (yellow) and the stimulus was subtracted from the data. **B**. Following artifact removal multi-unit action potentials were detected using a threshold detector (black line). Action potentials with characteristic spike profiles (inset) could be detected as early as 1 ms post-stimulation (red line). **C**. An exemplar channel shows a clear increase in multi-unit spikes 1-2 ms post-stimulus. **D**. To capture mesoscale connectivity the neural response in all electrodes within an array were pooled to reveal the global ICMS-evoked neural activity between sensorimotor regions (red trace). The significance of responses was assessed by randomly altering the temporal relationship between the ICMS pulse and neural activity across 1000 individual ICMS pulses (gray traces). In the exemplar shown, stimulation in the M1 distal forelimb region produced a strong short latency (<2.5 ms) increase in neural activity within PMd.

All multi-unit activity was aligned to the stimulus onsets (Fig. 1C). To determine the mesoscale ICMS-evoked firing for a single array, multi-unit spike times were pooled across each channel on an array and the global stimulus-evoked neural activity in the first 25 ms following the stimulus pulse was estimated using a kernel smoothing estimate with a normal kernel function (Fig. 1D). A reshuffling procedure was used to determine the neural activity expected by chance. For each stimulus pulse, a random start time within the inter-stimulus period was chosen. Multi-unit spike times were then aligned to this time point, pooled across channels with data from the beginning of the inter-stimulus period wrapped around to the end, and the chance global firing rate was estimated using a kernel smoothing estimate. This reshuffling procedure was repeated 10,000 times to produce estimates of the chance neural activity. To examine directly connected regions, we considered the transmission time of corticocortical connections. Based on estimates of corticocortical conduction velocity (33), and distances between electrode sites used (3-12 mm), we expected conduction delays of 0.33-3.0 ms for antidromic potentials with an additional synaptic delay for monosynaptic orthodromic potentials (34, 35). Accounting for these delays, the short-latency stimulus-evoked neural activity was estimated by summing the stimulus-evoked activity in the first 2.5 ms following the stimulus pulse. Statistical significance was determined by indexing the actual short-latency stimulus-evoked neural activity into the distribution of reshuffled neural activity with a significance threshold of p < 0.05 with Bonferroni correction for the total number of pairs of regions tested in each monkey. For significant connections, the depth of modulation was calculated by dividing the short-latency stimulus-evoked neural activity by the mean of the reshuffled data.

### Comparison with Anatomical Connectivity

Because these experiments were followed by an imaging experiment (28), it was not possible to use anatomical tracers to directly compare ICMS-evoked neural activity to anatomical connectivity. Therefore, ICMS-evoked neural activity was compared to literature-based anatomical connectivity (18-21, 30, 36-41). Studies using anterograde and retrograde anatomical tracer injections into the sensorimotor system in New World monkeys were examined to develop a map of sensorimotor connectivity (Fig. 2 and Table 1). Where quantitative estimates were available, a connection between two regions was classified as minor (present with < 2% of connections), moderate (2-10%), or major (> 10%). In studies without direct quantification, we reviewed and qualitatively assigned connectivity categories. Because connections with M1 are heterogeneous with specific connections into rostromedial (rm), rostrolateral (rl), and caudal (c) M1, we subdivided these regions within the map (20, 38, 41). Additionally, because it is uncertain if a unique area 2 is present in squirrel monkeys (31), we pooled reports of area 2 and area 5 connectivity for the region caudal to area 1. All connections were assumed to be reciprocal (18-21).

**Table 1:** Literature-reported anatomical connection strengths.

| Target Region | Connection Region | Connection Strength | Species | References |
| --- | --- | --- | --- | --- |
| **M1** | PMv | +++ | Squirrel Monkey, Owl Monkey, Cebus Monkey | (18-20, 37, 41) |
|  | PMd | +++ | Squirrel Monkey, Owl Monkey, Cebus Monkey | (19, 20, 37) |
|  | 3a | ++/+++ | Squirrel Monkeys, Owl Monkeys | (20, 37) |
|  | 3b | + / ++ | Squirrel Monkeys, Owl Monkey | (20, 37) |
|  | 1 | ++ | Squirrel Monkeys, Owl Monkey | (20, 37) |
|  | 2 | ++ | Owl Monkey | (20) |
| **PMv** | M1rl | +++ | Squirrel Monkey, Owl Monkey, Cebus Monkey | (18, 19, 21, 37, 38, 41) |
|  | 3a | + / ++ | Squirrel Monkey, Owl Monkey | (18, 21, 37) |
|  | 3b | + | Squirrel Monkey, Owl Monkey | (18, 21, 37) |
|  | 1 | + / ++ | Squirrel Monkey, Owl Monkeys | (18, 21, 37) |
|  | 2 | ++ | Owl Monkey | (21) |
| **PMd** | M1rm | +++ | Squirrel Monkey, Owl Monkey, Cebus Monkey | (19, 21, 37, 38) |
|  | 3a | - / ++ | Squirrel Monkey, Owl Monkey | (21, 37) |
|  | 3b | - / + | Squirrel Monkey, Owl Monkey | (21, 37) |
|  | 1 | + | Squirrel Monkey, Owl Monkey | (21, 37) |
|  | 2 | + | Owl Monkey | (21) |
| **3a** | PMv | + | Squirrel Monkey | (18) |
| **3b** | M1c | + / ++ | Squirrel Monkey, Titi Monkey | (30, 39, 40) |
|  | PMv | + | Squirrel Monkey | (18) |
|  | PMd | + | Squirrel Monkey | (39) |
| **1** | M1c | ++ | Squirrel Monkey, Titi Monkey | (30, 36) |
|  | PMv | + | Squirrel Monkey, Titi Monkey | (30, 36) |
|  | PMd | + | Squirrel Monkey, Titi Monkey | (30, 36) |
| **1/2** | PMv | + | Squirrel Monkey | (18) |
| **2/5** | M1 | ++ | Titi Monkey | (30) |
|  | PMv | ++ | Titi Monkey | (30) |
|  | PMd | ++ | Titi Monkey | (30) |

Connection Strength: +++ Major Connection, ++ Moderate Connection, + Minor Connection, - Absent Connection

**Figure 2:**
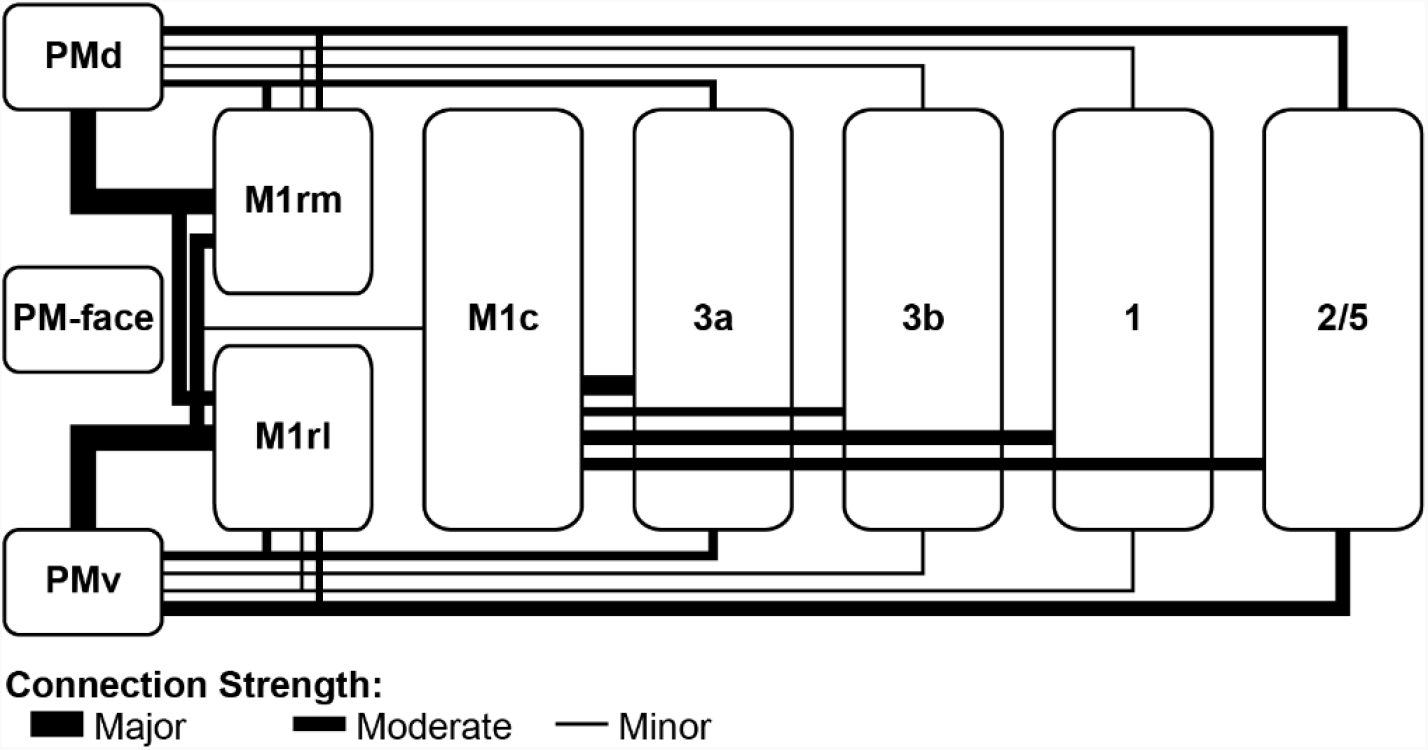
Anatomical connectivity map. The expected anatomical connectivity between sensorimotor regions of the squirrel monkey was compiled from prior literature sources using anatomical tracers to assess connectivity in New World non-human primate species (Table 1).

All array insertions were targeted to specific areas; however, the exact location was dependent upon the individual pattern of vasculature and extent of the craniectomy. To account for these differences in electrode locations between monkeys, the expected connectivity between electrodes was determined by weighting the sensorimotor connectivity with mixing matrices representing the exact location of each array. The current required to activate a region of cortex using ICMS is proportional to the square of the distance from the stimulating microelectrode, with estimates of the excitability constant ranging from 100-4000 μA/mm^2^ (13-15). Using our stimulus current of 50 μA the expected directly activated cortical region would have a radius of 0.1 – 0.7 mm. For this work, the stimulated region was modeled using a border of 0.5 mm around the stimulated microelectrode array. The connections from each microelectrode array were estimated by weighting the connections from each region by the percent of the stimulated area within each region. For the recorded electrode array, the connections to each region were weighted by the percentage of the electrode array within each region. To determine the correspondence between anatomical connectivity and short-latency ICMS-evoked activity, Pearson’s r was calculated between the pair-wise anatomical connectivity strengths and the depth-of-modulation of ICMS-evoked activity. Statistical significance of this correspondence was determined by rearranging the areal labels 10,000 times and recalculating the correlation coefficient. The p-value was determined by indexing the actual correlation coefficient into the surrogate distribution of correlation coefficients.

### Stability of ICMS-Evoked Neural Activity

Finally, we sought to determine whether ICMS-evoked neural activity was stable across the time course of the experiment. Because an array was always placed into M1, the premotor-M1 and S1-M1 connections were examined in multiple runs within each monkey. To determine whether the ICMS-evoked neural activity was stable within an animal across multiple testing sessions, the correlation (Pearson’s r) between the ICMS-evoked neural activity from a single stimulus-recording pair of regions in an individual recording and the ICMS-evoked neural activity from all other pairs of regions from all recordings was calculated. Correlation coefficients were grouped into repeated tests in which the correlation was between ICMS-evoked activity from repetitions within the same regions, and different comparisons in which ICMS-evoked activity between different pairs of regions were compared. The statistical significance for each monkey was evaluated using a Wilcoxon’s rank-sum test.

## Results

### ICMS-Evoked Neural Activity

After removing the stimulus artifact, multi-unit action potentials could be detected ∼1 ms post-stimulus with consistent spike waveforms (Fig. 1B). After aligning to the repeated single-pulse ICMS stimuli, clear increases in multi-unit firing could be observed within a few milliseconds (Fig. 1C). When pooled across all channels in an electrode array, the global ICMS-evoked neural activity showed short-latency responses with peaks in neural activity within 2.5 ms post-stimulus (Fig. 1D).

### Relationship between ICMS-evoked Neural Activity and Anatomy

In Monkey 1, stimulating/recording arrays were targeted to areas 1, 3b, 3a and M1, and a recording array was targeted to PMd, PMv, and PM-face. The 3a site overlapped 3a/3b border and the M1 array was placed into rostromedial M1 (Fig. 3A). Figure 3 compares the anatomical connectivity and ICMS-evoked neural activity. Based upon the anatomy, strong connections were expected between M1rm and PMd, and moderate connections were expected between M1rm and PMv. Significant increases in ICMS-evoked neural activity were observed for both pairs of regions with the strongest response in PMd following M1rm stimulation. Across all connections there was a strong correlation between the strength of the expected anatomical connectivity and the ICMS-evoked response (Pearson’s r = 0.79, p = 0.0023).

**Figure 3:**
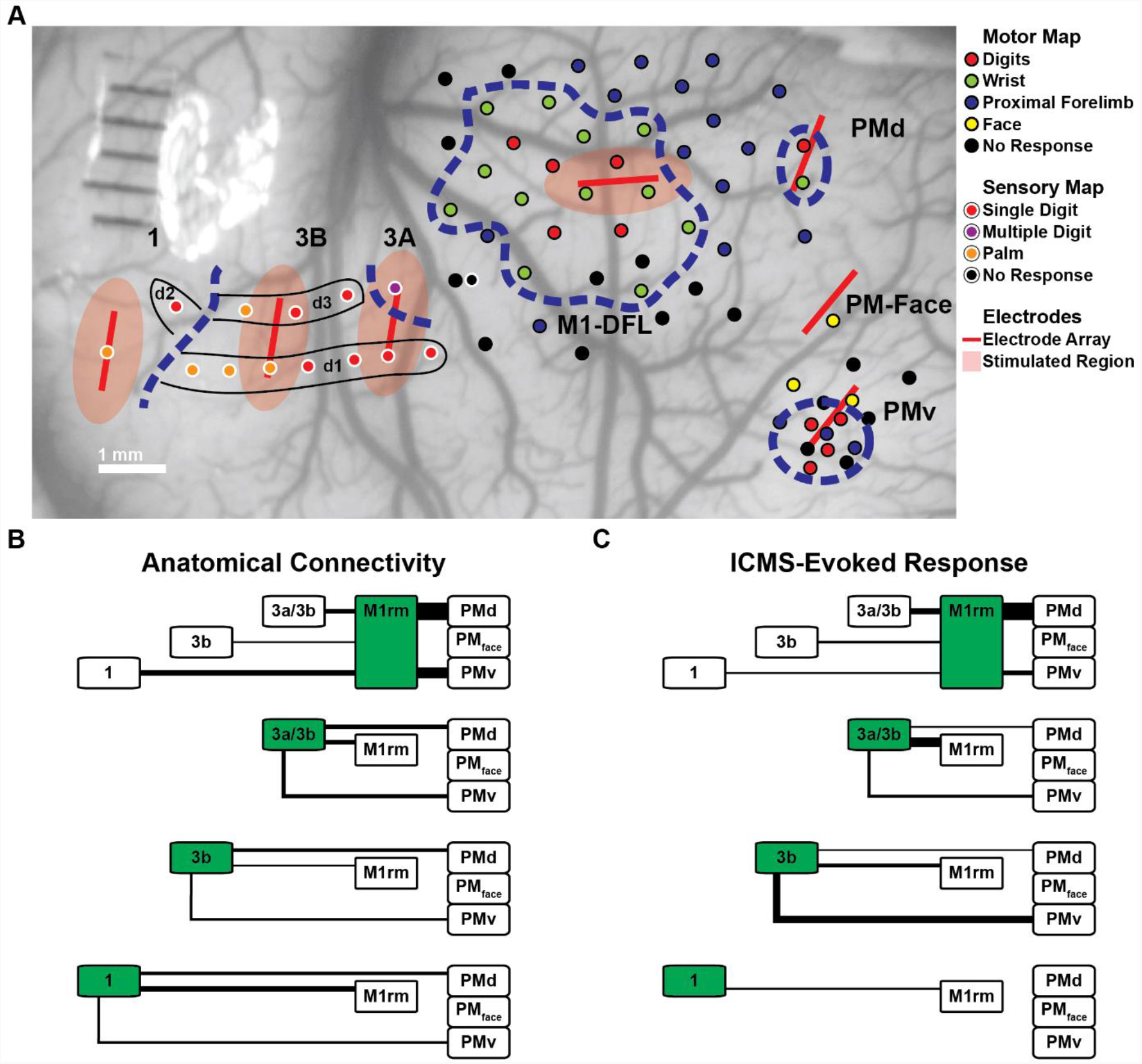
Comparison between anatomical and ICMS-evoked connectivity (Monkey 1). **A**. In Monkey 1 stimulating/recording arrays were placed in the rostromedial portion of the M1 distal forelimb region, across the area 3a/3b border, in area 3b, and area 1, and recording arrays were placed in PMd, PMV, and the face region of M1 in between PMD and PMv. Motor regions were assessed using ICMS mapping (black-outlined sites), and sensory region boundaries were assessed using sensory mapping (white-outlined sites). Arrays (red lines) were targeted to single brain regions, but the specific placement was adjusted to fit the vasculature pattern. Red shadings show the regions expected to be activated by the 50 μA ICMS pulse **B**. Expected anatomical connectivity was assessed by weighting the literature-based anatomical connectivity map by the portion of each anatomical region expected to be stimulated and recorded from using the specific electrode locations and regional boundaries (**A**). Line thicknesses represent the normalized number of anatomical connections **C**. Short-latency ICMS-evoked neural activity was evaluated for each connection. Regions highlighted in green represent stimulated areas and the line thicknesses show the depth-of-modulation of ICMS-evoked increases in neural activity. The ICMS-evoked activity was significantly correlated to the expected anatomical connectivity (r = 0.79, p = 0.0023), with the strongest connectivity observed between M1rm and PMd.

In Monkey 2, stimulating/recording arrays were targeted to areas 2/5, 1, 3b, 3a, M1, PMd, PMv, and PM-face. Each region was recorded, and stimulation was performed in all regions except PM-face. The area 2/5 array overlapped area 1 and the M1 array was placed into rostrolateral M1. Additionally, the area stimulated by the electrodes in areas 1 and 3a likely overlapped with area 3b. Figure 4 shows a comparison of the anatomical connections and ICMS-evoked neural activity. Based upon the anatomy, strong connections were expected between M1rl and PMv, and moderate connections were expected for the M1rl-PMd, M1rl-3a, 3a-PMv, and 2/5/1-PMv connections. Significant increases in ICMS-evoked neural activity were observed for each of these connections except the 2/5/1-PMv connection with the strongest response in PMv following M1rl stimulation. Additionally, ICMS-evoked responses were observed in the PM-face region following stimulation in M1rl and area 2/5/1 but not in response to the other 3 stimulated regions. Across all connections there was a moderately strong correlation between the strength of the expected anatomical connectivity and ICMS-evoked response (Pearson’s r = 0.61, p = 0.0003).

**Figure 4:**
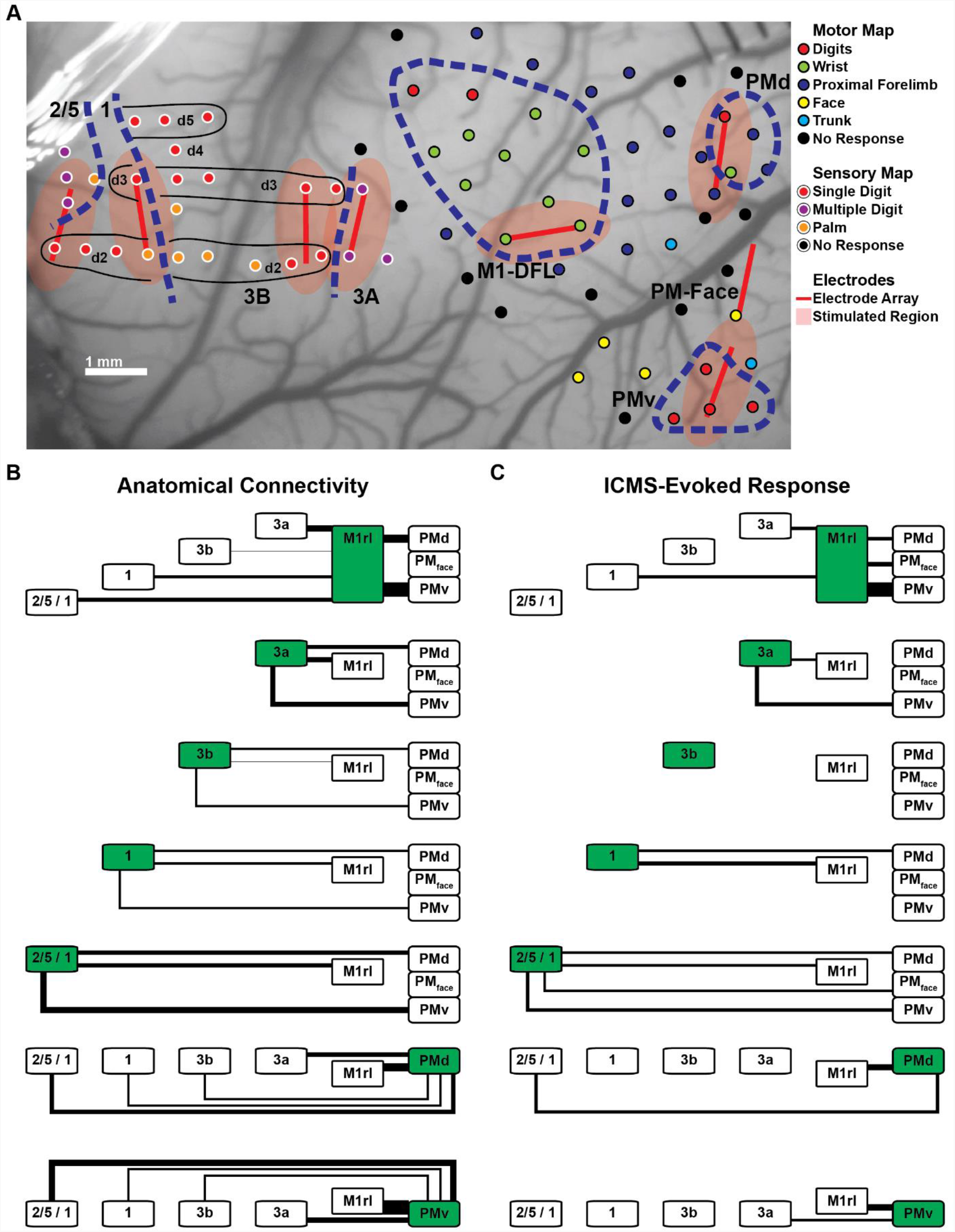
Comparison between anatomical and ICMS-evoked connectivity (Monkey 2). **A**. In Monkey 2 stimulating/recording electrodes were placed in the rostrolateral portion of the M1 distal forelimb region, in PMv, PMd, the face region of premotor cortex, area 3a, area 3b, area 1, and across the border between area 1 and area 2/5. All regions were stimulated except for PM-face. The specific placement of arrays (red lines) was adjusted to fit the vasculature pattern. Red shadings show the area expected to be activated by the 50 μA ICMS pulse **B**. Expected anatomical connectivity was assessed by weighting the literature-based anatomical connectivity by the portion of specific regions stimulated and recorded from (**A**). Line thicknesses represent the expected number of anatomical connections **C**. Short latency ICMS-evoked neural activity was evaluated for each connection. Regions highlighted in green represent stimulated areas and the line thicknesses show the depth-of-modulation of ICMS-evoked increases in neural activity. The ICMS-evoked activity was significantly correlated to the expected anatomical connectivity (r = 0.61, p = 0.0003), with the strongest connectivity observed between M1rl and PMv.

In Monkey 3, stimulating/recording electrodes were targeted to areas 1, 3b, 3a, M1, PMd, and PMv. The M1 electrode was placed into M1rm and stimulation in area 1 likely activated a small portion of 3b. Figure 5 shows a comparison of the anatomical connectivity and ICMS-evoked neural activity. Based upon the anatomy, moderate to strong connections were expected between M1rm and PMd, PMv, and 3a, and between 3a and both PMd and PMv. Significant increases in ICMS-evoked neural activity were observed for each of these connections with the strongest response for the bidirectional M1rm-PMd and M1rm-PMv connections. Across all connections there was a strong correlation between the strength of the expected anatomical connectivity and ICMS-evoked response (Pearson’s r = 0.87, p < 0.0001).

**Figure 5:**
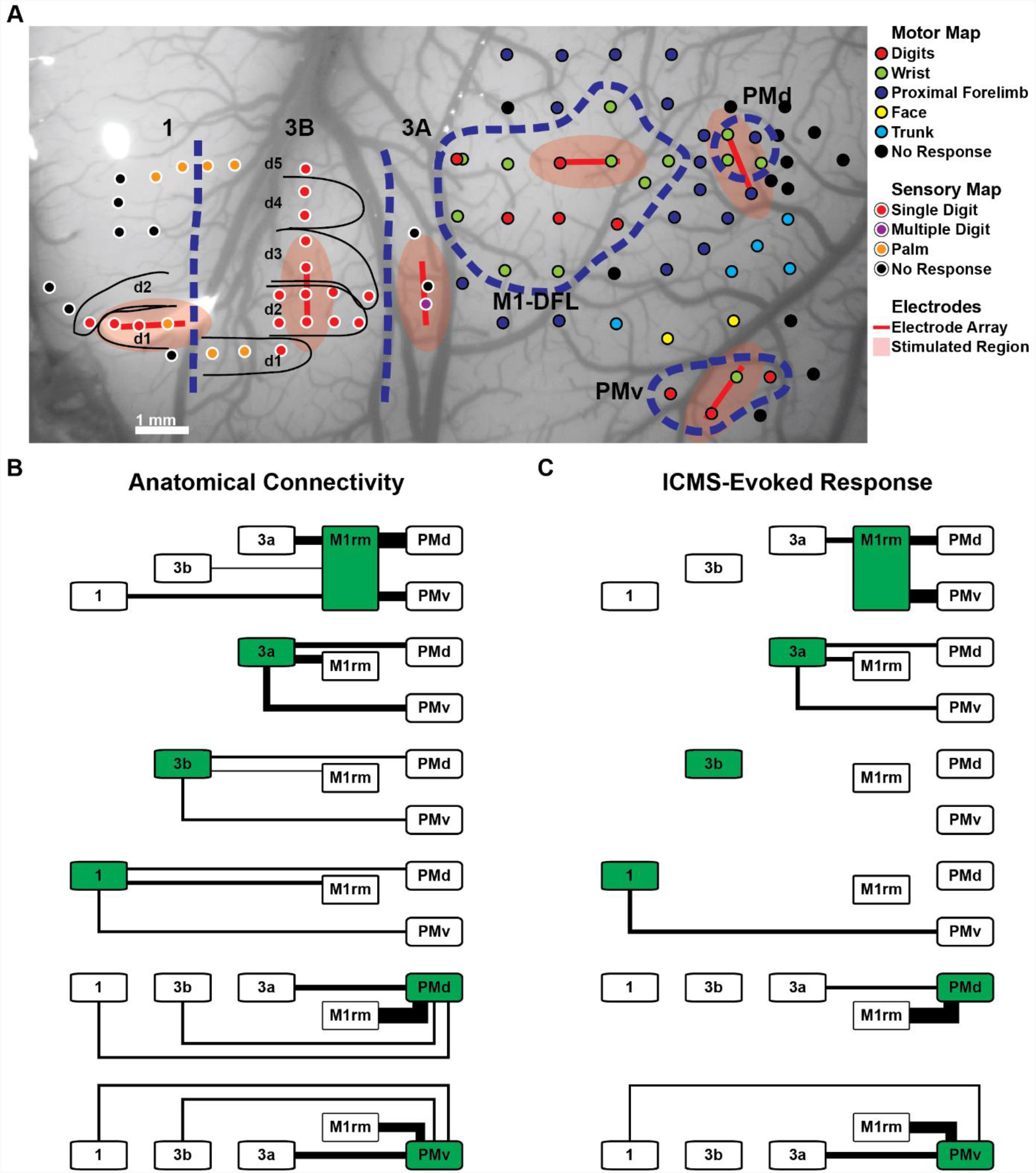
Comparison between anatomical and ICMS-evoked connectivity (Monkey 3). **A**. In Monkey 3 stimulating/recording arrays were placed in the rostromedial portion of the M1 distal forelimb region, PMv, PMd, area 3a, area 3b, and area 1. Motor regions were assessed using ICMS mapping (black-outlined sites), and sensory region boundaries were assessed using sensory mapping (white-outlined sites). The specific array placement was adjusted to fit the vasculature pattern. Red shadings show the regions expected to be activated by the 50 μA ICMS pulse **B**. Expected anatomical connectivity was assessed by weighting the literature-based anatomical connectivity by the portion of each region stimulated and recorded from. Line thicknesses represent the expected number of anatomical connections **C**. Short latency ICMS-evoked neural activity was evaluated for each connection. Regions highlighted in green represent stimulated areas and the line thicknesses show the depth-of-modulation of ICMS-evoked increases in neural activity. The ICMS-evoked activity was significantly correlated to the expected anatomical connectivity (r = 0.87, p < 0.0001), with significant ICMS-evoked neural activity in the regions expected to have moderate-strong anatomical connectivity (3a, M1, PMd, and PMv).

Taken together, the known anatomical connections strongly correlated with ICMS-evoked neural activity in each monkey. Figure 6 shows a confusion matrix comparing the strength of anatomical connections to the strength of ICMS-evoked activity across monkeys. The correspondence is driven by the separability of regions with moderate-major anatomical connections and regions with no known anatomical connections. All of the major connections demonstrated moderate or major modulation of neural activity following ICMS, and 16 of 17 connections with moderate anatomical connectivity showed statistically significant ICMS-evoked increases in neural activity. Finally, 7 of 9 connections with no anatomical connections showed an absence of ICMS-evoked neural activity.

**Figure 6:**
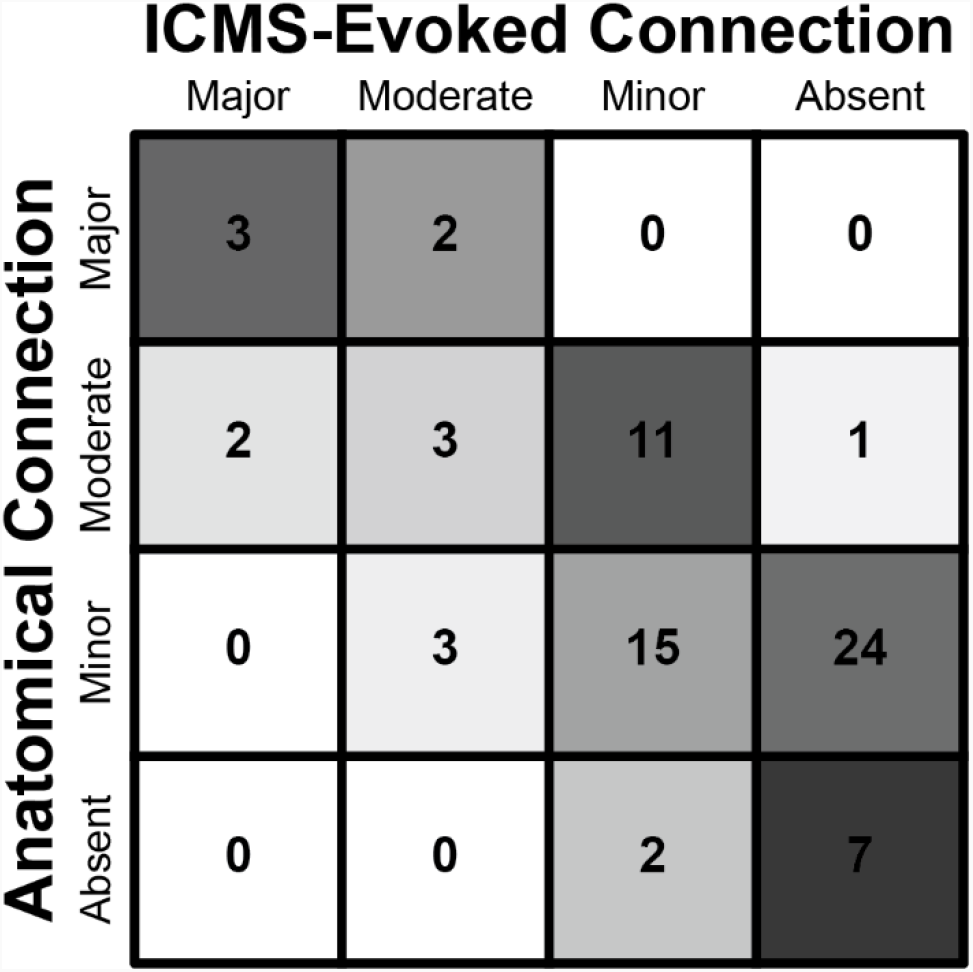
Anatomical and ICMS-evoked connectivity confusion matrix. The strength of anatomical and ICMS-evoked activity was linearly discretized and compared. The numbers within the confusion matrix shows the number of connections with specific levels of anatomical and ICMS-Evoked connection strength. Colors were normalized to the total number of connections in each anatomical category with darker colors indicating more connections. Across monkeys, the ICMS-evoked activity correlated with anatomical connectivity. This effect was driven by the presence of significant ICMS-evoked neural activity when moderate-major anatomical connectivity is present, and an absence of ICMS-evoked neural activity when no anatomical connectivity has been reported. Shadings are normalized to the total number of connections in each category of anatomical connectivity.

### Stability of ICMS-Evoked Responses

Because the PM and S1 electrodes were moved sequentially, connections between M1 and both PM and S1 regions were examined 2-4 times in each monkey. Figure 7A shows exemplar patterns of ICMS-evoked neural activity for each repetition of two pairs of electrode positions. The amplitude and time course of ICMS-evoked neural firing was maintained across recordings. Across all pairs of regions in each monkey, the correlation between ICMS-evoked neural activity from different recordings was higher for repeated electrode positions than different electrode positions (Fig 7B) (Monkey 1: p < 0.0001; Monkey 2: p < 0.0001; Monkey 3: p = 0.0004).

**Figure 7:**
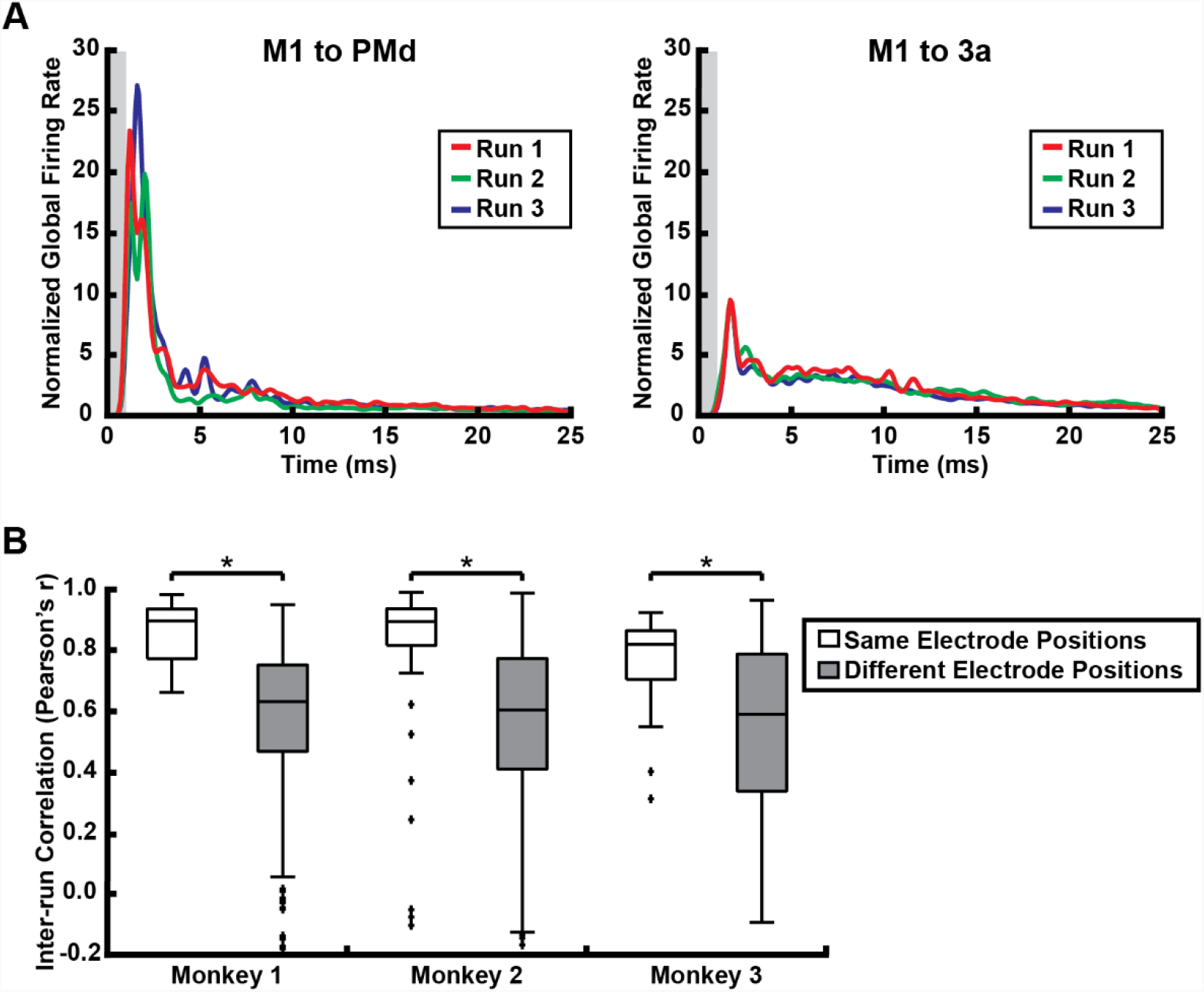
Stability of ICMS-evoked neural responses. To examine the stability of ICMS-evoked neural activity, connections that were evaluated in multiple recordings were examined. **A**. Exemplar responses are shown from Monkey 1 in PMd (left) and area 3a (right) after M1 stimulation. Each area shows a clear and distinct stimulus-evoked increase in neural activity that was maintained across recordings. **B**. The stability across connections was evaluated by comparing the correlations of ICMS-evoked neural activity between pairs of recordings with repeated electrode positions to the correlations of ICMS-evoked neural activity between pairs of recordings with different electrode positions. The ICMS-evoked responses for repeated electrode positions were significantly more correlated than for different locations, showing the ICMS-evoked neural responses were stable over the course of the experiment in each monkey.

## Discussion

These results show that the observed strength of ICMS-evoked neural activity is significantly correlated to previously described anatomical connectivity. While the horizontal spread of neural activity within one electrode array has previously been examined (42), this is the first demonstration of the correspondence between mesoscale anatomical and evoked connectivity using ICMS-evoked neural activity. This effect was driven primarily by the presence of ICMS-evoked neural activity between regions with major and moderate anatomical connections. There was more variability in the regions with sparse anatomical connections. Because of the limited number of expected anatomical connections between these regions, it is likely that the placement of any specific microelectrode location has a comparatively greater impact on the pattern of ICMS-evoked activity. Despite this potential limitation, short-latency ICMS-evoked neural activity appears to be a strong surrogate for effective anatomical connectivity, therefore we believe it is appropriate to refer to this technique as “Stimulation-Evoked Effective Connectivity” (SEEC).

The specific mechanisms that underlie SEEC are an important consideration when comparing it to anatomical connectivity. Because the volume of axons activated and the density of activated soma increases monotonically with stimulus intensity, it is thought that the dominant mode of ICMS is likely activation of axons (12). However, low amplitude stimulation can also lead to transsynaptic activation (16). Therefore, by considering the neural activity <2.5 ms after each ICMS pulse, a combination of antidromic action potentials and monosynaptic orthodromic action potentials were likely measured. While it is possible to isolate antidromic responses by timing stimulation to naturally occurring action potentials to test for collisions between orthodromic and antidromic potentials (13, 33), this method is impractical across the large number of brain regions tested with multiple sites per array and multiple units detected on many electrode sites. However, because anatomic connectivity between the regions studied is reciprocal (18-21), it is unlikely that the results were impacted by this mixture of antidromic and orthodromic responses.

When pairs of regions were examined multiple times, SEEC was stable over the duration of the experiment, despite slight shifts in electrode positions. Therefore, this measure is invariant to slight changes in anesthetic state and minor changes in the neural population sampled. While these results do not directly address the stability of ICMS-evoked connectivity over longer periods, their within-session repeatability is a minimum criterion for chronic monitoring and assessment. There are likely several important considerations that may impact the transition from acute to chronic electrodes. First, these experiments were done under ketamine anesthesia, which potentially disrupts baseline corticocortical information transfer (43), therefore increased background levels of shared corticocortical neural activity in awake animals may limit SEEC measurements between weakly connected regions. Second, the acute arrays used here sampled neural activity at multiple sites within each cortical column. Chronic arrays could sample from a wider spatial region, leading to less dependence on the specific neural population sampled. Finally, the encapsulation of chronic electrodes could impact the ability to drive stimulation and record neural activity, however as ICMS thresholds for sensory perception decrease over time (5) and we utilize multi-unit firing, it is unlikely that the ability to measure ICMS-evoked responses would be significantly impacted.

Several other methods have also been proposed to examine corticocortical connectivity. In particular, ICMS has also been used in conjunction with fMRI or intrinsic optical imaging to measure intra- and interregional connectivity (22-26, 44). Furthermore, similar to our findings, ICMS-evoked neural activity measured with intrinsic optical imaging corresponds with both anatomical connectivity and resting-state functional connectivity (45). However, the stimulation amplitudes in these studies were several times higher than the amplitudes used here, increasing the volume of direct activation, and potentially the risk of cortical damage with repeated stimulation (46). Additionally, while an area of cortex can be imaged repeatably, fMRI in animal models requires the use of anesthesia for scanning and it is technically challenging to maintain the visual field necessary for repeated optical imaging. Finally, because these measures capture the hemodynamic response, it is uncertain whether they represent subthreshold synaptic potentials or action potentials (26). Importantly, applying our SEEC method in chronic electrode arrays would allow us to examine longitudinal changes in direct effective connectivity in awake animals.

The ability to perform longitudinal experiments using chronic arrays is particularly significant for studies examining recovery from neural injuries. Following a stroke, neuroplasticity leads to changes in connectivity within and between unaffected cortical regions. Within the motor system, these alterations can be seen in changes in motor map outputs (7, 9), alterations in patterns of functional connectivity (47-50), and alterations in anatomical connections (51). Because specific changes in map output and connectivity have been observed after functional recovery (7, 9), it is thought that there may be a causal relationship. However, as the time course of motor map reorganization is distinct from the time course of behavioral recovery (8), the specific relationship between changes in connectivity and behavioral recovery remains uncertain. Because causal relationships must demonstrate temporal precedence, determining the specific relevance of changes in corticocortical connectivity with respect to behavioral changes will require assessing changes in connectivity with high temporal specificity.

There are several limitations to note with the current approach. Because the animals were used for a subsequent experiment, we were unable to compare SEEC to anatomical connections within the same animals. However, the literature-based estimates of anatomical connections, combined with cortical mapping to determine specific electrode locations, likely produced accurate estimates of anatomical connectivity between sites at the mesoscale level. Second, the experiments used acute microelectrode arrays that were repeatedly reinserted and could possibly damage the cortex. However, the stability of SEEC shows that any local damage to neural populations did not significantly impact the results. Finally, the stimulation paradigm used single-pulse ICMS delivered at 200 ms intervals. This constant interstimulus interval could lead to some entrainment of neural activity (52). However, changes in neuronal excitability and entrainment should impact the reshuffled control data in addition to the ICMS-evoked response. Therefore, it is unlikely that the stimulus timing impacted the results presented.

## Conclusion

We have demonstrated that neural activity evoked from ICMS can be used to examine mesoscale corticocortical connectivity. This will be especially valuable for examining longitudinal changes in corticocortical connectivity associated with a brain injury, neural disorder, or therapeutic interventions. Future studies will extend these analyses to chronic experiments in healthy animals and animals recovering from cortical lesions.

